# Chicken cecal microbial functional capacity and resistome differ by age and barn disinfection practice

**DOI:** 10.1101/2024.05.23.595585

**Authors:** Yi Fan, Tingting Ju, Tulika Bhardwaj, Douglas R. Korver, Benjamin P. Willing

## Abstract

Chemical disinfectants and water-wash methods are widely employed in sanitizing broiler chicken barns. Previous studies showed that chemical disinfectants affect environmental microbial composition and antibiotic resistance genes (ARGs). However, little is known regarding how barn disinfection treatments impact the chicken gut resistome and microbial functionality. The current study compared the effects of chemical disinfection and water-wash method on the gut microbiome and resistome of commercial broilers using a crossover experimental design after 2 production cycles at 7 barns. Shotgun metagenomic sequencing performed on cecal contents collected at day 7 and 30 also allowed evaluation of age-associated characteristics of microbiome. Age of the chickens had the largest effects on the resistome, with younger birds having increased relative abundance of total ARGs (P<0.05) and differences in resistance mechanism, however, functional and resistome differences were also identified by barn sanitation practice. At day 7, chickens in chemically-disinfected barns had decreased functional capacity related to amino acid synthesis compared to the water-wash group. Additionally, genes related to stringent response were enriched in chickens raised under chemically-disinfected condition (FDR-P<0.05), suggesting selection for stress resistance. Consistently, lower abundance of genetic pathways encoding amino acid biosynthesis associated with cecal *Helicobacter pullorum* was observed in the disinfection group at day 30 compared to the water-wash group, with the same pattern in short-chain fatty acid biosynthesis (FDR-P<0.05). Overall, while the use of disinfectants in barn sanitation slightly affected the relative abundance of some ARGs in the gut, age had a dominant effect on the microbial functionality and resistome.

**Importance:** This is the first study to evaluate the effect of sanitation practices on microbial functional capacity and resistome of chickens in a commercial setting. It is also amongst the biggest metagenomics studies on the gut microbiome of broiler chickens. It provides new insights into the changes in resistance profiles with age that agree with other studies examining maturation of the microbiome in other species. Finally, the current study provides valuable insights for informing industry sanitation practices and future studies on broiler gut microbiome and resistome.

## Introduction

Livestock farming accounts for over 50% of antibiotic usage globally (1). Compared with other livestock species, chickens were reported to have the highest density of antibiotic resistance genes (ARGs) due to the high stocking density and short production cycle (2). Currently, biocidal agents such as benzalkonium chloride (BAC), hydrogen peroxide, glutaraldehyde, ethanol, and sodium hypochlorite have been widely applied in agriculture for disinfection purposes (3). While chemical disinfectants can inhibit antibiotic-resistant bacteria and destruct ARGs through oxidation, they may also induce bacterial adaptation, potentially through promoting antibiotic resistance through co-selection (4). The impact of chemical disinfectants on ARG proliferation remains uncertain, with some studies suggesting that they may act as stressors, stimulating the proliferation and transfer of microbial ARGs (5–7). For example, BAC, a commonly used quaternary ammonium compound, has been associated with increased resistance to ampicillin, cefotaxime, and sulfamethoxazole in various food-related bacterial isolates (5, 8), along with co-selected ARGs (9). Some studies, on the other hand, suggested that chemical disinfectants may contribute to controlling antibiotic resistance by reducing the abundance of ARGs. For instance, quaternary ammonium compounds and sodium hypochlorite used in treating swine manure have been reported to decrease the abundance of selected ARGs (*erm(B), erm(C), erm(F), intI1, tet(Q),* and *tet(X)*) (10). Oxidants like chlorine and hydroxyl radicals have also demonstrated potential in eradicating ARGs presented in both *E. coli* cells and plasmids(11). With the controversial effect of chemical disinfectants on ARGs, there is limited information available regarding the effects of chemical disinfectant-treated rearing environments on the resistome in the gut of animals. In this sense, barn cleaning practices, which involve chemical disinfectants, may impact the presence and persistence of ARGs throughout production cycles. Consequently, from the perspective of food safety and environmental sustainability, it is important to explore the influence of chemical disinfectants usage in barn cleaning.

In broiler chicken production, barn sanitation has been used with the goal of enhancing biosecurity and prevent disease transmission between flocks. Both full sanitation with chemical disinfectants (FD) and water-wash (WW) method are widely employed in the Canadian poultry production system (12) with FD being required on a yearly basis. Our previous study showed that FD resulted in an undesired increased carriage of *Campylobacter jejuni* accompanied by alterations in the cecal microbial composition, revealed by 16S rRNA gene amplicon sequencing (13). Full disinfection also resulted in lower cecal short-chain fatty acids (SCFAs) when compared to the WW (13). In the current study, we therefore sought to gain greater insight into the effects of barn sanitation and age on the functionality of the gut microbiome, specifically in terms of microbial metabolic capacity and the profile of antibiotic resistance genes.

## Materials and methods

### Animals

The current study was performed according to the guidelines of the Canadian Council on Animal Care with approval of the University of Alberta Animal Care and Use Committee (AUP00002377). Broiler chicken management and barn cleaning practices were described previously (13). Briefly, the animal study was conducted on commercial broiler chicken farms in Alberta, Canada. During each production cycle, samples were collected from barns that had undergone two consecutive rounds of repeated cleaning treatments, including both FD and WW. For FD treatment, manure and litter were completely removed from the barn after chickens were depopulated. Subsequently, chemical disinfection was performed using foam containing 7% sodium hydroxide, 7% 2-(2-2-butoxyethoxy) ethanol, 6% sodium laureth sulfate, 5% sodium N-lauroyl sarcosinate, and 5% tetrasodium ethylenediaminetetraacetic acid on all surfaces within the facilities, followed by high- and low-pressure water rinse with water temperature set at 35℃. After the facilities were air-dried, foam containing 10% glutaraldehyde, 10% benzalkonium chloride, and 5% formic acid was applied to surfaces of the facilities for 60 mins followed by high-pressure water rinse, overnight air-dry, and fresh litter placement. For WW treatment, manure and used litter were removed, followed by low-pressure water rinse with the water temperature set at 35℃ for all facility surfaces, air-dry overnight, and fresh litter placement.

A cross-over design was performed which resulted in a total of 14 production flocks with 7 flocks assigning to each treatment (FD or WW). The study included a total of 140 chickens with 35 chickens sampled from each treatment at each age (day 7 or day 30). In each production flock, Ross 308 broiler chicks were placed within 12 h post-hatch and confined to half of the house, then allowed access to the entire house starting at D7. The flock size was approximately 14,000 birds, with a final stocking density of 30 kg/m^2^. All chickens were fed an antibiotic free diet *ad libitum* and sent for processing at 32-35 days of age when the average target live weight of 1.8 kg was reached. At D7 and D30 of age, five broilers per flock were randomly selected from different areas within each barn and euthanized using cervical dislocation. Approximately 300 mg of cecal contents were collected using sterile technique, placed on dry ice until being transported to the lab, and stored at -80 ℃.

### Shotgun metagenomic sequencing

The total DNA extraction process was conducted as previously outlined (13). In summary, DNA extraction was carried out from homogenized litter samples and cecal contents using the QIAamp Fast DNA Stool Mini Kit (Qiagen, Valencia, CA, USA), including an extra bead-beating step utilizing approximately 200 mg of garnet beads at a speed of 6.0 m/s for 60 seconds (FastPrep-24 5G instrument; MP Biomedicals Inc., Santa Ana, CA, USA). Library preparation and shotgun sequencing were performed at the Genome Quebec Innovation Centre (Montreal, Canada). Libraries for all samples were prepared using the Nextera™ DNA Flex Library Prep Kit (Illumina Inc., San Diego, CA, USA), and the subsequent shotgun sequencing was executed on a NovaSeq 6000 system (Illumina Inc., San Diego, CA, USA). Read quality was assessed using FastP v0.23.2, and low quality reads, sliding windows, adaptors, polyG, and duplicated sequences were removed (14). Kneaddata v0.10.0 was used to remove host DNA contaminants (https://github.com/biobakery/kneaddata). Briefly, a chicken host reference database was built using bowtie2 v2.4.1 with genome *Gallus_gallus* 105 release from Ensembl (15). Subsequently, reads aligned to the host genome were removed as host contaminants. Microbial taxonomic classification was profiled using kraken2 (v2.1.2) (16), and the relative abundance estimation was conducted by Bracken2 (v2.6) (17) Bacterial taxa appeared less than 5% of the samples were filtered out for subsequent analyses. Genome assembly was performed via megahit (v1.2.9) with default parameters (18). The abundance of functional genes and enriched pathways were estimated using HuMAnN3 (v3.0.1) based on the UniProt 90 database followed by annotation using the Metacyc database (19, 20). The relative abundance of aligned genes and pathways were normalized to copy numbers per million reads using the HuMAnN3 utility scripts (20).

Antibiotic resistance-encoding genes were annotated against the Comprehensive Antibiotic Resistance Database (CARD, version 3.1.4) via Resistance Gene Identifier (RGI, version 5.1.0) with a cut-off set at 95% identity (21). To reveal the distributions of microbial taxa, functional genes, and ARGs across samples, principal coordinate analysis (PCoA) based on Bray-curtis distance metric was performed via vegan package in R (version 3.6.1). To evaluate dispersion, the R package “betadisper” was used to calculate distance to centroid. Due to potential biases introduced by variations in the relative abundance of bacterial taxa, 16S rRNA copy numbers harbored by different bacterial species as well as fluctuation caused by ARGs located on mobile genetic elements, identified ARGs per sample were normalized to reads per million total ARG reads for subsequent comparisons.

### Statistical analyses

Permutational Multivariate Analysis of Variance (PERMANOVA) tests were used to determine clustering significance using the adonis function in the vegan package in R (version 3.6.1, false discovery rate (FDR) adjusted *P* < 0.05). Differentially abundant microbial taxa, gene pathways and ARGs associating with treatments or age were identified using LDA effect size (LEfSe) implemented in the lefser R package (Bioconductor version 3.15) with a significant cut off set at LDA score > 2 and FDR adjusted *P* < 0.05. Differential abundant analysis of profiled gene pathways and ARGs between groups was performed using DESeq2 package in R (Bioconductor version 3.15) (22). Significance of differential abundance was determined with FDR adjusted *P* < 0.05 and log2 fold change > 1. Except for the statistical analyses mentioned above, GraphPad Prism 8 (Graphpad Software, San Diego, CA, USA) was used to conduct statistical analyses. To determine differential significance of the microbiome alpha diversity indices, the Kruskal–Wallis test was used with significance set at *P* < 0.05. Two-way ANOVA was used to assess total ARG reads between different barn sanitations practices and sampling timepoints. The Spearman correlation was used to correlate ARG abundance and the relative abundance of bacterial species. Correlation significance was determined by the corr.test function with FDR adjusted *P* < 0.05, and a moderate association was determined by ∣R∣ > 0.4. Correlation was visualized using the corrplot package in R (version 3.6.1).

## Data availability

The sequences from the current study have been submitted to the NCBI Sequences Read Archive under the BioProject ID: PRJNA1108021.

## Results

### Cecal microbial structures and functional capacities were impacted by both barn cleaning methods and age

In this study, shotgun metagenomic sequencing was used to provide a comprehensive profile of the cecal microbial community to further assess the effects of barn cleaning methods. An average of 52,133,725.06 ± 1,265,988.36 quality-controlled reads per sample (mean ± SEM) were obtained and processed using kraken2, resulting in 34,252,401.09 ± 6,541,062.17 reads per sample aligning to the RefSeq bacteria database. At the phylum level, Firmicutes, Bacteroidetes, Proteobacteria, and Actinobacteria made up the majority cecal microbial communities both at day 7 and 30. However, there were shifts in the relative abundance of these phyla from D7-to D30-microbiota (D7 vs. D30: Firmicutes, 62.14% vs. 24.59%; Bacteroidetes, 25.57% vs. 53.27%; Proteobacteria, 8.21% vs. 15.59%; Actinobacteria, 0.76% vs. 5.92%, other phyla, 3.32% vs. 0.63%). PERMONOVA analyses based on Bray-Curtis distance dissimilarity matrix revealed that barn cleaning methods had limited impact (adonis *P* = 0.23, Figure 1a) on the D7 microbial community structure and a modest impact on D30 microbiota (R^2^ = 0.02, adonis *P* < 0.01, Figure 1b). However, there were distinct separations between D7 to D30 cecal microbiota irrespective of barn cleaning method (Figure 1c, adonis *P* < 0.001). The D30 microbiota had greater distance to the centroid compared with the D7 microbiota (distance to centroid = 0.48 and 0.56 for D7 and D30, respectively, FDR adjusted *P* < 0.01), indicating that the D7 microbiota had greater homogeneity. In addition, the Shannon index indicated that barn cleaning methods had minimal impact on alpha diversity at D7 (*P* = 0.96) and D30 (*P* = 0.25); however, differences were observed between D7 and D30 microbial communities as reflected by increased Shannon diversity with age (*P* < 0.05) (Figure 1 d-f). In terms of differentially abundant taxa, LEfSe analysis revealed no taxa with differences in the relative abundance between FD and WW groups at D7 (FDR adjusted *P* > 0.05). However, at D30, *Ruminococcus torques, Faecalibacterium prausnitzii, Barnesiella viscericola*, and *Helicobacter pullorum* were enriched in the WW group, whereas *Megamonas funiformis* was higher in the FD group (FDR adjusted *P* < 0.05, LDA >2, Figure 2). In addition, successional changes on the cecal microbial composition were demonstrated by LEfSe analysis. Briefly, 11 and 8 bacterial families were associated with the D7 and D30 chicken cecal microbiota, respectively (Figure S1).

**Figure 1.**
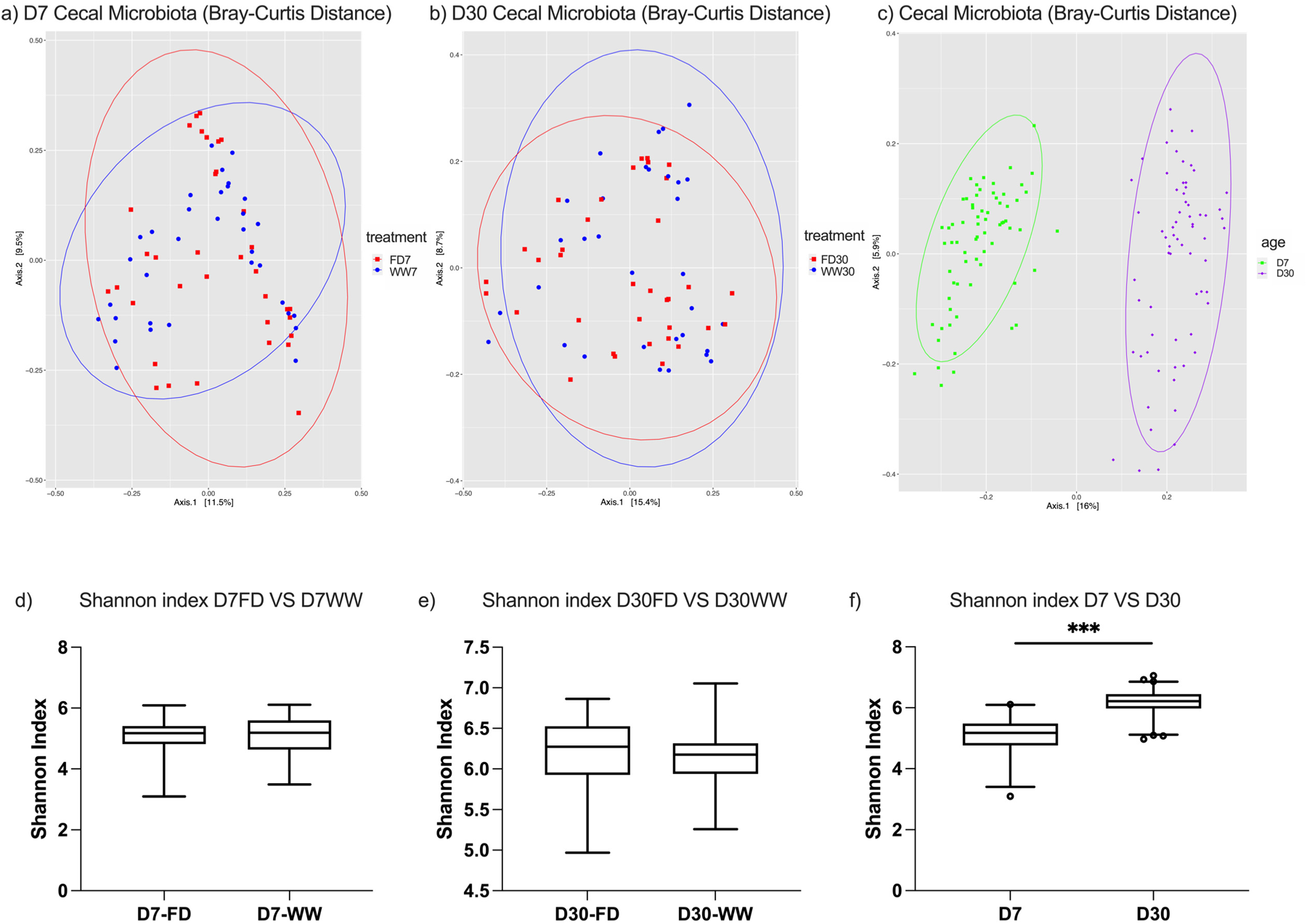
Broiler chicken cecal microbial structure affected by the barn cleaning practices and age. Figure a-c. Factors impacting the cecal microbial structures. Age had a major impact on microbial compositions, whereas the cleaning methods had a modest effect on the D30 cecal microbiota. Figure d-f. Factors affecting the cecal microbial alpha-diversity as indicated by the Shannon index. At D30, the richness and evenness of the cecal microbial species significantly increased compared to that at D7. FD, full disinfection; WW, water-wash; ***, *P* < 0.001.

**Figure 2.**
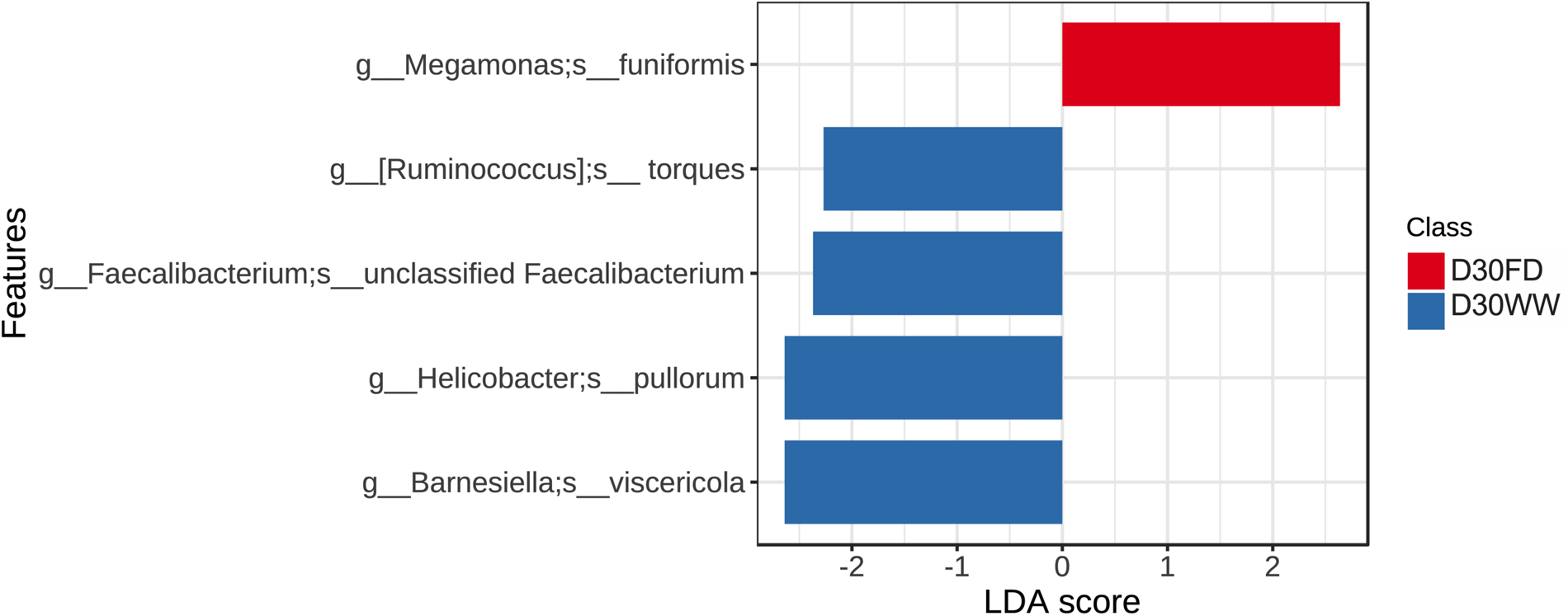
Differential abundant bacterial species between barn cleaning practices at day 30 suggested by LEfSe analysis (LDA score > 2 and FDR adjusted *P* < 0.05). At day 30, *Ruminococcus torques, Barnesiella viscericola, Helicobacter pullorum*, *Faecalibacterium prausnitzii* were more abundant in the ceca of the chickens from the WW treatment group, whereas *Megamonas funiformis* was more abundant in the chicken cecal microbiota of FD group. FD, full disinfection; WW, water-wash.

Cecal microbial functionalities were subsequently profiled and annotated, resulting in an average of 64.63% ± 1.18% of the total reads per sample (mean ± SEM) being mapped to the UniProt 90 database and further annotated. At D7, six pathways were altered by the barn cleaning methods (Figure 3a, log2 fold-change > 1, FDR adjusted *P* < 0.05), including the sucrose degradation pathway IV (PWY-5384), the L-cysteine biosynthesis pathway VI (PWY-I9), the super-pathway of UDP-glucose-derived O-antigen building blocks biosynthesis (PWY-7328), the UDP-N-acetyl-D-glucosamine biosynthesis pathway (UDPNAGSYN-PWY), the phospholipase pathway (LIPASYN-PWY), and the stringent response guanosine 3’-diphosphate 5’-diphosphate metabolism pathway (PPGPPMET-PWY). Among these altered pathways, PWY-5384 was mainly harbored by *Escherichia coli, Lactobacillus* spp., and *Bifidobacterium* spp. At D30, a series of pathways were altered by barn cleaning methods (Figure 3b). Specifically, twelve pathways were enriched in the WW group, including the pyruvate fermentation to acetate and lactate pathway II (PWY-5100), the L-lysine biosynthesis pathway I (DAPLYSINESYN-PWY), the L-lysine biosynthesis pathway II (PWY-2941), the L-isoleucine biosynthesis pathway I (ILEUSYN-PWY), the L-methionine biosynthesis pathway III (HSERMETANA-PWY), the L-isoleucine biosynthesis pathway III (PWY-5103), the phosphatidylglycerol biosynthesis pathway I (PWY4FS-7), the phosphatidylglycerol biosynthesis pathway II (PWY4FS-8), the ADP-L-glycero- and β-D-manno-heptose biosynthesis pathway (PWY0-1241), the CMP-3-deoxy-D-manno-octulosonate biosynthesis pathway (PWY-1269), the super-pathway of phospholipid biosynthesis I (PHOSLIPSYN-PWY), and the super-pathway of branched chain amino acid biosynthesis (BRANCHED-CHAIN-AA-SYN-PWY), whereas the FD group had an enriched pathway involved in hexitol fermentation to lactate, formate, ethanol and acetate (P461-PWY) (Figure 3b). The total of 13 altered pathways at D30 were attributed by 62 different bacteria species. The enriched PWY-5100 in the WW cecal microbial community could be considered as a consequence of an increased level of *H. pullorum* (Figure 4, FDR adjusted *P* < 0.05, LDA >2), whereas the contribution of *Lachnoclostridium* sp. *An76* was relatively smaller (FDR adjusted *P* < 0.05, LDA = 1.36). Indeed, the observed pathway enrichment linked to amino acid syntheses at the WW group were mainly attributed to *H. pullorum*. Compared with the modest impact on cecal microbial functional capacities by barn cleaning methods, age was shown as a strong factor affecting cecal microbial functionalities. A total of 89 pathways were significantly different between D7 and D30 cecal microbiome (Table S1, log2 fold-change > 1, FDR adjusted P < 0.05, DESeq2 analysis with sanitation treatments as blocking factor).

**Figure 3.**
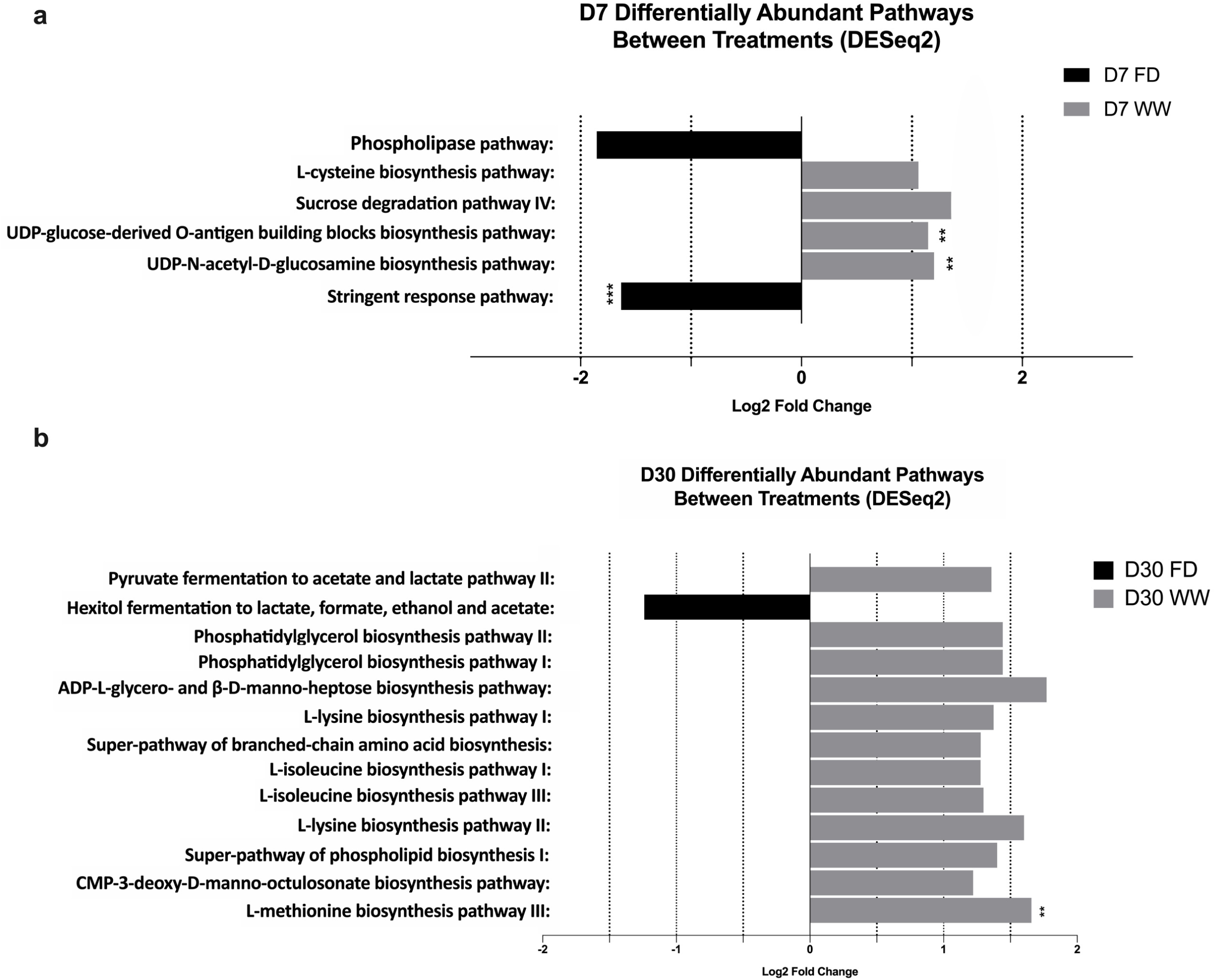
Microbial functional pathways that were significantly impacted by the barn cleaning treatments at day 7 (a) and day 30 (b) revealed by DESeq2. The graph shows differentially abundant genetic pathways harbored by the chicken cecal microbial communities at day 7 and day 30 suggested by DESeq2, respectively (FDR *P* < 0.05, log2 fold-change >1). a) At day 7, the FD-derived chicken gut microbiome had enriched stringent response pathway coupled with decreased abundance of pathways linked to amino acid synthesis, saccharide degradation, and bacterial cell wall synthesis. b) At day 30, the FD-derived chicken gut microbial functional capacity had decreased abundance of genetic pathways linked to multiple amino acid syntheses. D7, day7; D30, day 30; FD, full disinfection; WW, water-wash **, FDR *P* < 0.01; ***, FDR *P* < 0.001.

**Figure 4.**
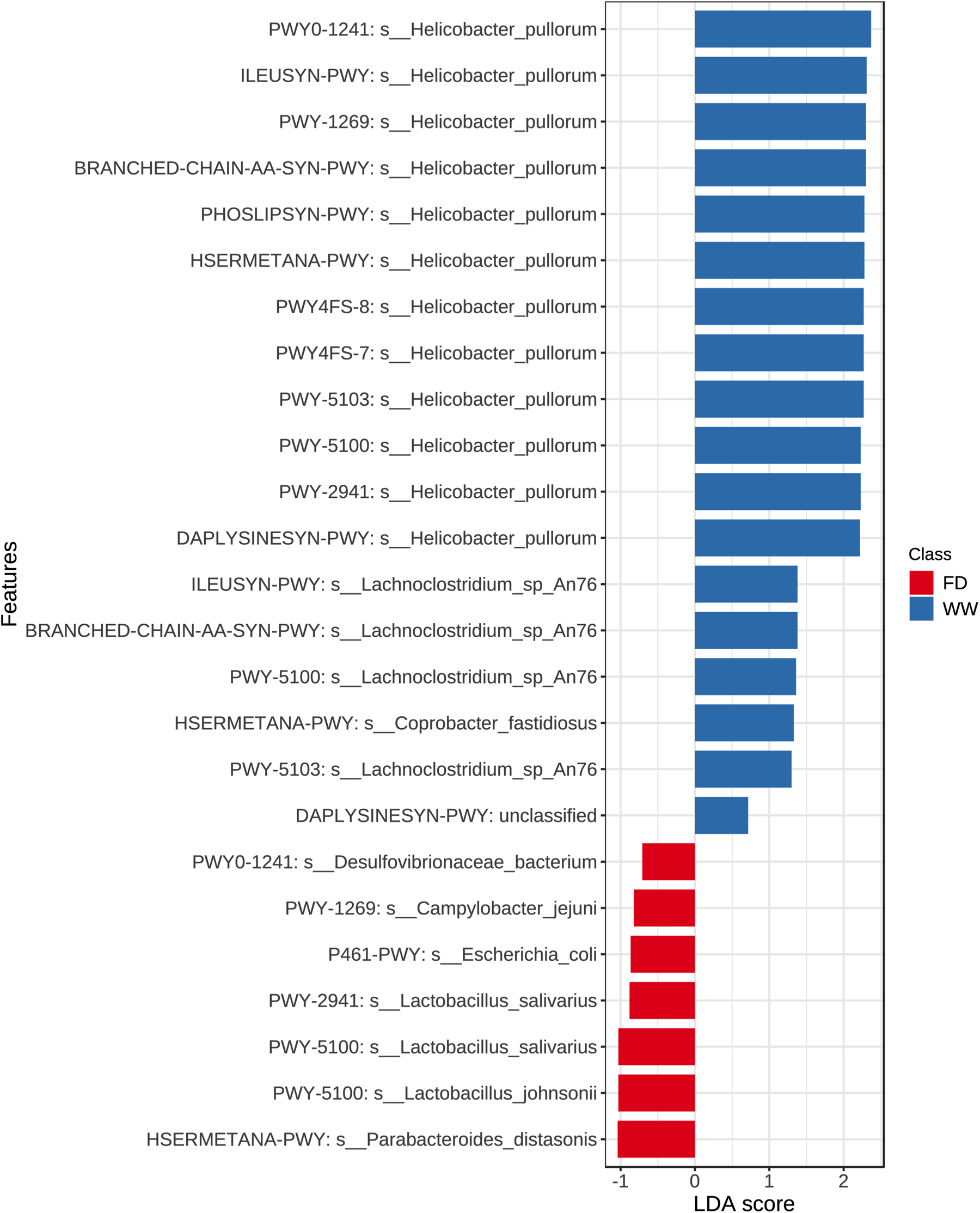
LEfSe results of metabolic pathways harbored by specific bacterial species and the association with treatments at day 30 (FDR adjusted *P* < 0.05). LEfSe result suggested that the increased abundant pathways in the WW group were mainly contributed by *Helicobacter pullorum.* FD, full disinfection; WW, water-wash.

### The effect of barn cleaning practices and age on the cecal resistome

An average of 0.133% ± 0.012% of total reads per sample (mean ± SEM) were mapped to the CARD database. Both barn cleaning methods and age had an impact on the resistome (Figure 5b, FDR *P* <0.05). Beta-dispersion analyses revealed that the cecal resistome of the 7-day old broiler chickens had greater distance to centroid compared to that in D30 broiler chickens (*P* < 0.01), indicating more variations in cecal antimicrobial resistance at D7 (Figure 5c). The relative abundance of total ARGs was higher at D7 compared with D30 (Figure 5a). Overall, a total of 496 ARGs from 60 gene families and 386 ARGs from 52 gene families were identified from D7 and D30-chicken cecal microbiome, respectively. Among the detected ARGs, the tetracycline-resistant gene (*tet*) *tetW* was most abundant, followed by lincosamide nucleotidyltransferase (*lnu*) *lnuC*, *tet*(*44*), *tet(W/N/W)*, *tetQ*, the erythromycin ribosomal methylation 23S ribosomal RNA methyltransferase (*erm*) *ermB*, aminoglycoside resistance gene (*APH*) *APH*(*3*)*-IIIa*, *tetO*, and *tet32.* In terms of gene families, tetracycline-resistant ribosomal protection proteins (RPPs) family was most abundant followed by the lincosamide nucleotidyltransferase (LNU, lincosamide-resistant) family and the major facilitator superfamily (MFS) antibiotic efflux pump (multi-drug resistant) family. These ARG families collectively accounted over 90% of the broiler chicken cecal resistome (92.08% ± 11.03% mean ± SEM). At D7, the most abundant ARG families in the cecal resistomes were the tetracycline-resistant RPP family, the MFS antibiotic efflux pump family, the resistance-nodulation-cell division (RND) antibiotic efflux pump (multi-drug resistant), the *erm* gene family, and the LNU family. Whereas at D30, the top 5 abundant gene families were the tetracycline-resistant RPP followed by the LNU family, the MFS antibiotic efflux pump family, the aminoglycoside nucleotidyltransferase (*ant*) family *ant*(*6*) (aminoglycoside-resistant), and the *erm* gene family. When characterizing the resistance mechanism and distribution of the ARGs, we discovered that antibiotic target protection, which mainly consisted of tetracycline resistance via tetracycline-resistant RPPs, was the most common mechanism of antibiotic resistance. In addition, antibiotic efflux and antibiotic inactivation were the second most common resistance mechanism found at D7 and D30, respectively (Figure 6). LEfSe results supported the successional change of the chicken cecal resistome (Figure 7, FDR *P* < 0.05, LDA > 2). In accordance with the different antibiotic mechanisms between D7 and D30, LEfSe analysis identified that 11 ARG gene families were associated with the age of the broiler chickens (Figure 7). Particularly, the MFS antibiotic efflux pumps and the RND antibiotic efflux pump showed the highest LDA score at D7, indicating that antibiotic efflux plays a key role in the early life cecal resistome. Whereas on D30, AMR gene families conferring antibiotic inactivation, such as the LNU family and the tetracycline inactivation enzyme, had increased predominance.

**Figure 5.**
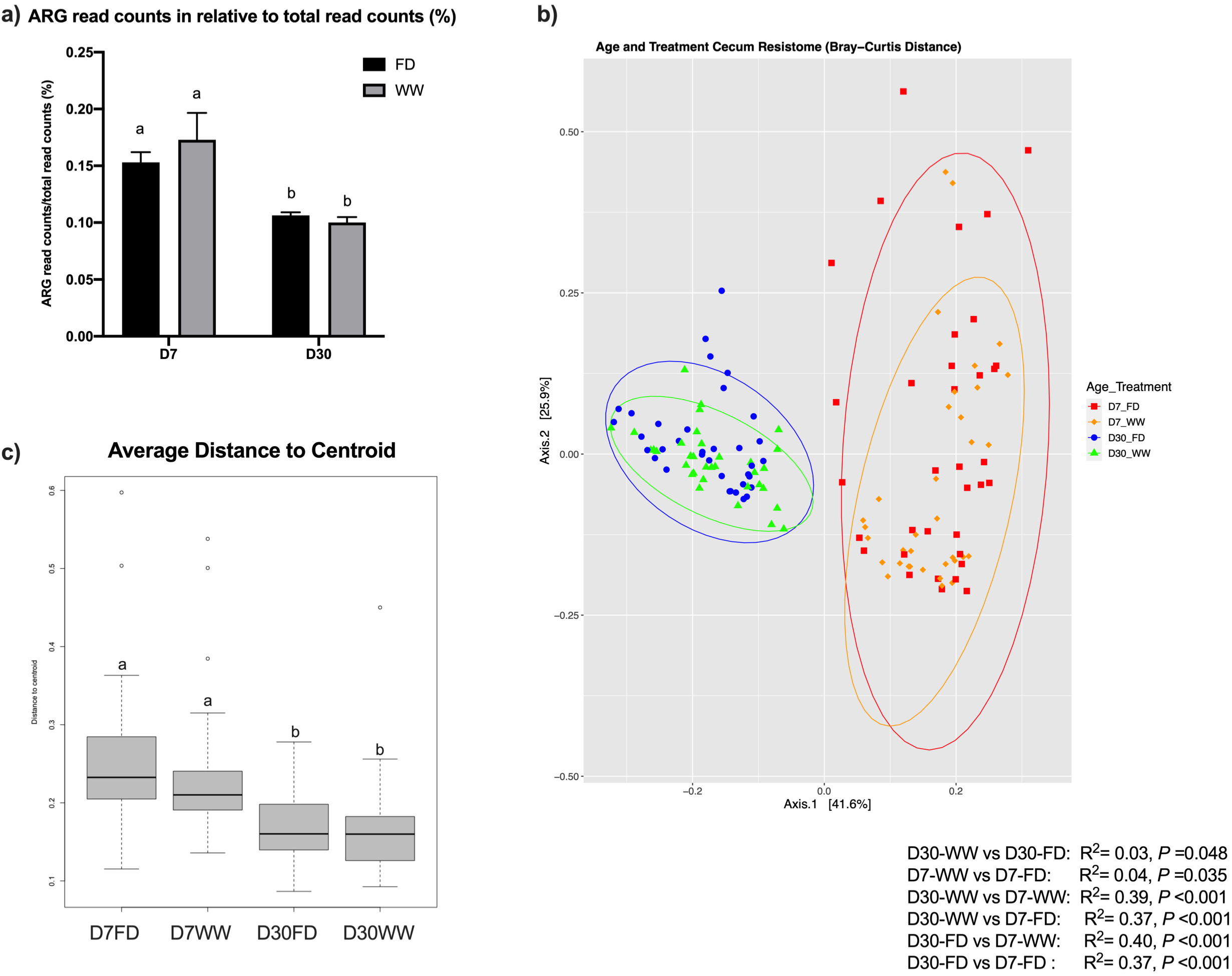
The chicken cecal resistome was affected by the barn sanitation practices and sampling timepoints. a) ARG read counts relative to total read counts. Generally, at D7, a higher percentage of ARG read counts was detected in the chicken cecal microbiota compared to D30. b) PCoA plots based on Bray-Curtis similarity distance matrix showing resistome clusters by treatments or age. Barn cleaning methods had modest effect on the distribution of microbial resistome. Age was the main driver of the resistome pattern. c) The average distance to centroid based on beta-disperse showed that the variation between resistomes was greater among the D7 chickens in comparison to the D30 chickens. ARG, antibiotic resistant genes; D7, day7; D30, day 30; FD, full disinfection; WW, water-wash.

**Figure 6.**
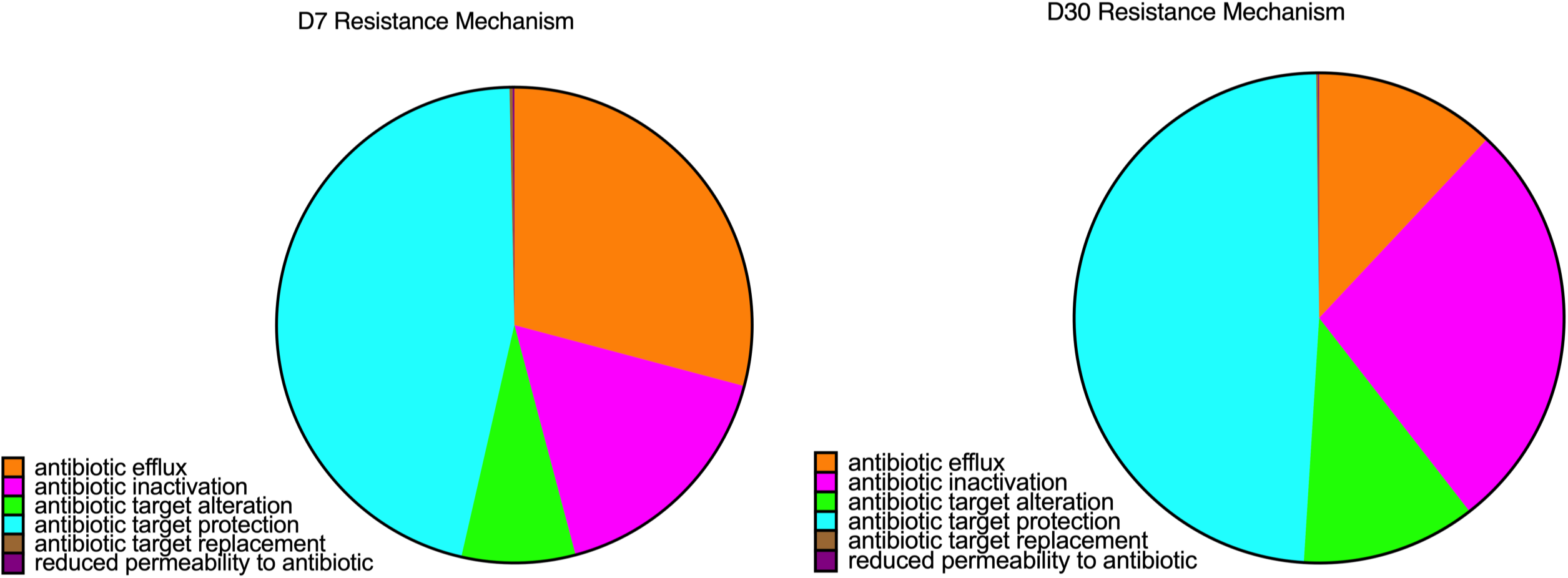
Antibiotic resistance mechanism conferred by detected ARGs in the cecal microbial resistomes of D7 and D30 chickens. Differences on the major antibiotic resistant mechanisms harbored by the 7-day and 30-day chicken cecal microbial resistomes were observed. With the mechanism of antibiotic target alteration being dominant on both ages, genes conferring antibiotic efflux and antibiotic inactivation were representative of the day 7 and day 30 chicken cecal resistome, respectively.

**Figure 7.**
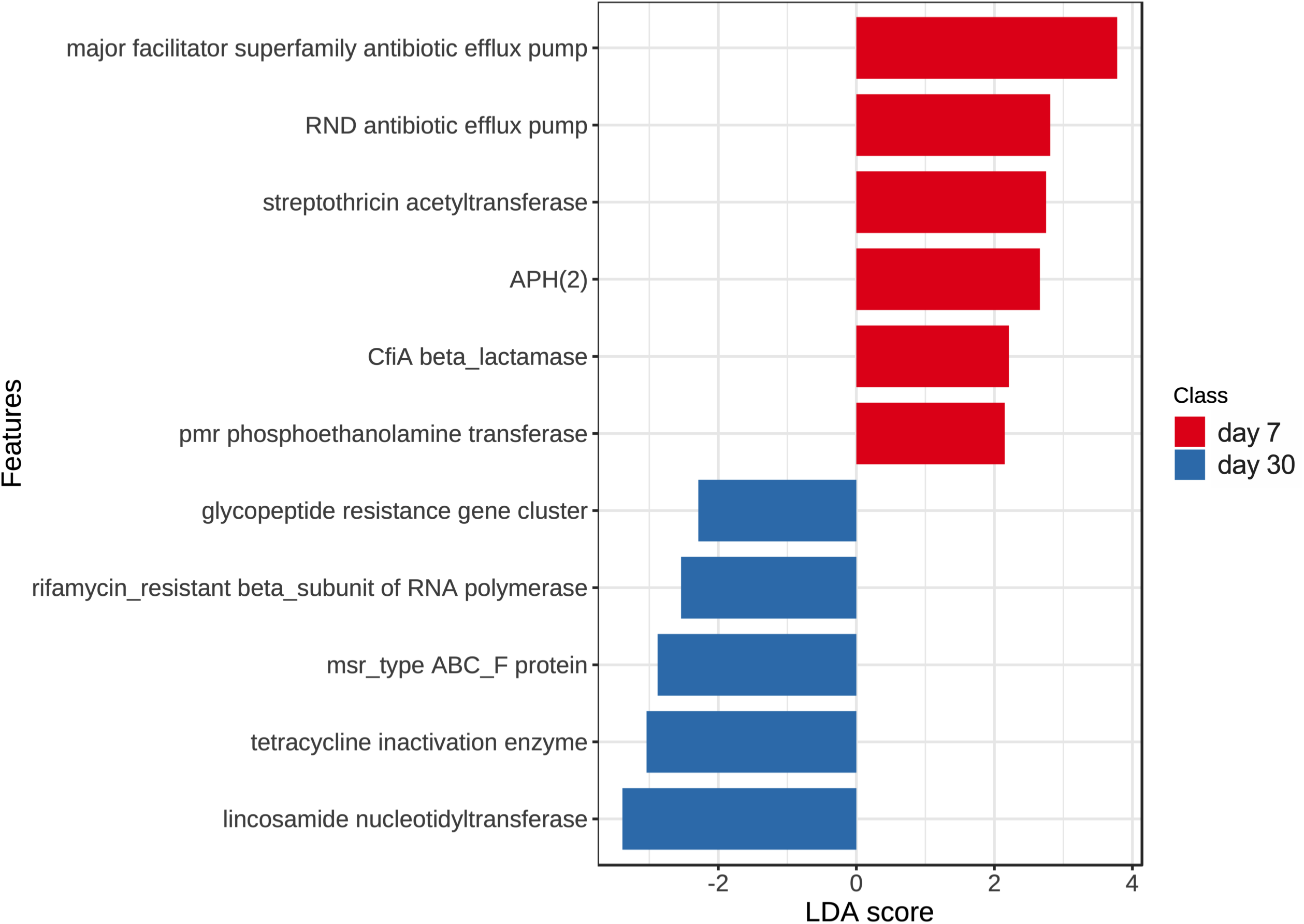
Differentially abundant cecal microbial antibiotic resistant gene families between day 7 and day 30 suggested by LEfSe. Graph showing antibiotic gene families associated with different ages (FDR *P* < 0.05, LDA > 2). At day 7, genes encoding antibiotic efflux pumps were representative of the chicken cecal microbial resistome. At day 30, antibiotic resistant genes conferring antibiotic inactivation (e.g. the lincosamide nucleotidyltransferases and tetracycline inactivation enzymes) were more predominant. RND, resistance-nodulation-cell division; APH, aminoglycoside resistance gene; pmr, polymyxin resistance; msr, macrolide resistance; ABC, ATP binding cassette.

Compared with the age effect, barn cleaning methods had a modest impact on the chicken resistome profile. Specifically, five AMR gene families including the *erm* gene family, rifamycin resistant beta-subunit of RNA polymerase (*rpoB*), streptogramin vat acetyltransferase, ATP-binding cassette (ABC) - F subfamily RPPs (macrolide- and lincosamide-resistant), and *vanR* glycopeptide resistance gene cluster were found to be more abundant in the D7-WW treatment compared with the D7-FD treatment (Figure 8, FDR *P* < 0.05). At the gene level, *cprR* (peptide antibiotic-resistant)*, ermB* (macrolide-, lincosamide- and streptogramin-resistant, MLS-resistant)*, lnuA* and *lnuB* (lincosamide-resistant)*, lsaE* (pleuromutilin-, streptogramin-, and lincosamide-resistant)*, oleB* (macrolide-resistant)*, tet(L)* and *tetM* (tetracycline-resistant)*, vatE* (streptogramin-resistant)*, Vibrio anguillarum* chloramphenicol acetyltransferase gene (phenicol-resistant), the *Bifidobacterium bifidum ileS* (mupirocin-resistant), the *Bifidobacterium adolescentis rpoB* (rifampicin-resistant), and *vanR* variants in the *vanA*, *vanG*, and *vanL* clusters (vancomycin-resistant) were more abundant in the D7-WW group. At D30, the impacts of cleaning methods on the chicken gut microbiome were subtle, indicating that the barn cleaning practices had greater impacts in younger chickens. No differentially abundant ARG gene families were identified between FD and WW by DESeq2. However, genes including *ermG* (MLS-resistant) and *vanR* variant in *vanI* cluster (vancomycin-resistant) were enriched in the D30-WW group (Figure 9).

**Figure 8.**
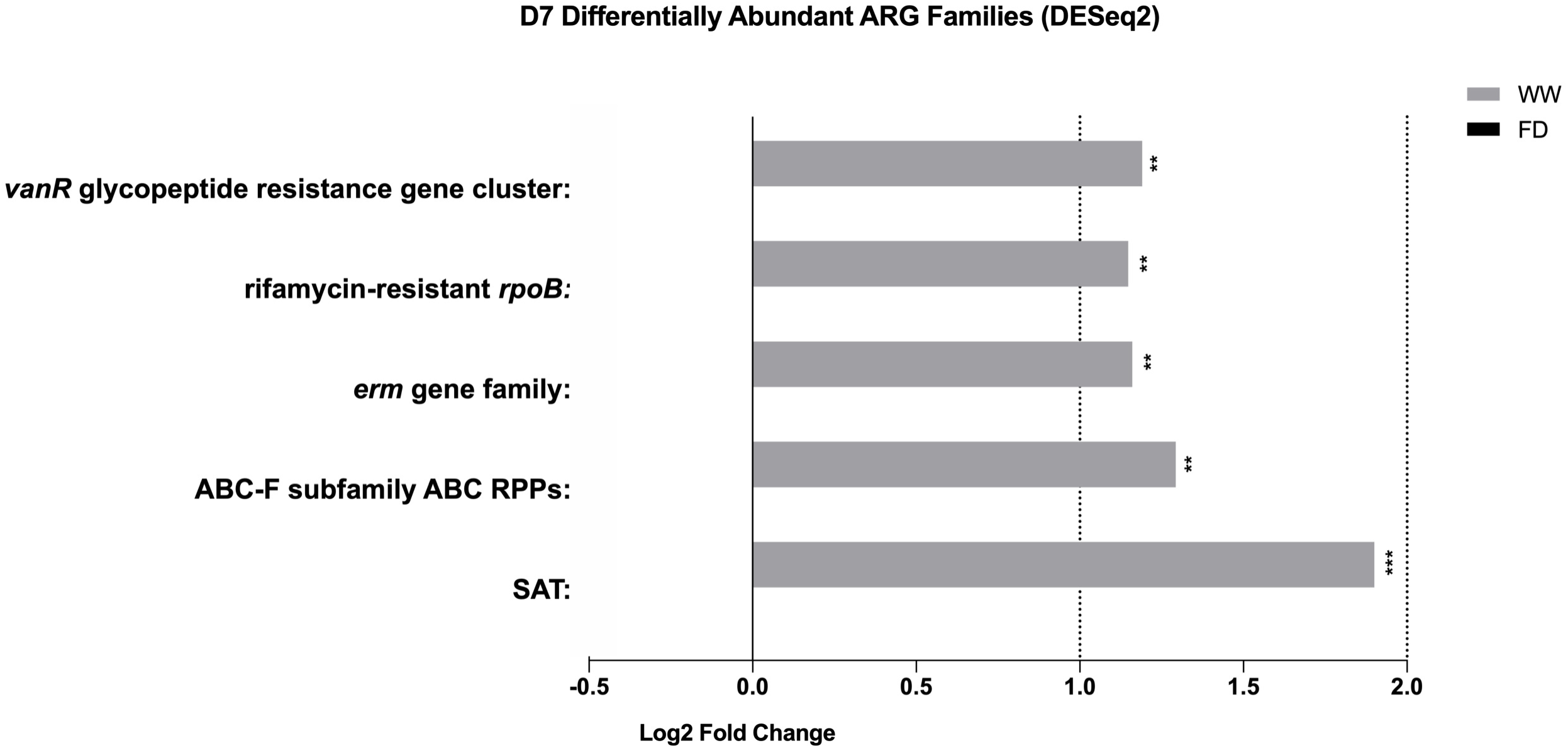
Differentially abundant antibiotic resistant gene families between FD and WW at D7 suggested by DESeq2. Graph shows differentially abundant antibiotic resistant gene families between barn sanitation practices at day 7 suggested by DESeq2 (FDR *P* < 0.05, Log2 fold change > 1). Some persistent ARG gene families (e.g. *erm* gene family) were enriched in the WW-derived chicken cecal microbiome at day 7. ARG, antibiotic resistant gene; *vanR*, vancomycin resistant gene R component; *rpoB,* gene encoding β-subunit of bacterial RNA polymerase; *erm,* 23S ribosomal RNA methyltransferase; ABC-F subfamily ABC RPPs, ABC-F ATP-binding cassette ribosomal protection protein genes; SAT, Streptogramin A acetyltransferase genes, FD, Full disinfection; WW, Water-wash, ***, *P* < 0.001; **, *P* < 0.01.

**Figure 9.**
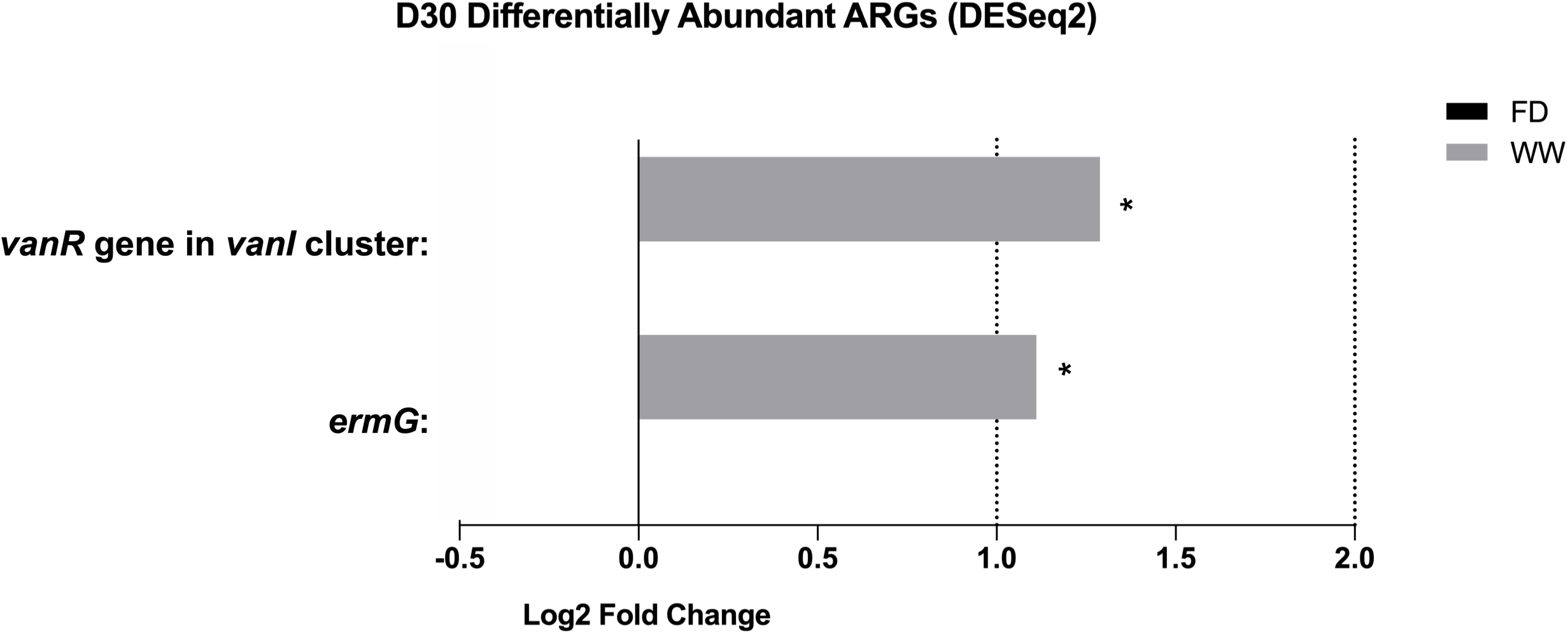
Differentially abundant antibiotic resistant gene between FD and WW at D30 suggested by DESeq2. Graph shows differentially abundant antibiotic resistant genes between barn sanitation practices at day 30 suggested by DESeq2 (FDR *P* < 0.05, Log2 fold change > 1). Compared to the impact of barn cleaning practices at day 7, the effects on the 30-day chicken gut microbial resistome was relatively smaller. On the gene level, *ermG* and *vanR* gene in the *vanI* cluster were enriched by the WW treatment. ARG, antibiotic resistant gene; *vanR*, vancomycin resistant gene; *erm,* 23S ribosomal RNA methyltransferase genes; FD, Full disinfection; WW, Water-wash, *, *P* < 0.05.

## Discussion

To date, limited information is available regarding how barn cleaning methods affect the chicken cecal microbiota. Using 16S rRNA sequencing technique, we previously reported that at D30, the genus *Helicobacter* was enriched in the WW group (13). In the current study, functional genetics analyses suggested that the barn cleaning methods altered the functional capacities of the chicken cecal microbiota. The exposure to FD at 1 day of life may impact the assembly of the early cecal microbiota of chicks, and thereby led to altered microbial functionalities observed later in life, which possess decreased genetic potential for amino acid and SCFA synthesis. In addition, we confirmed that chicken cecal *H. pullorum* harboring genes linked to SCFA and amino acid production was higher in the WW treatment.

Shotgun metagenomic sequencing of the samples provided taxonomic information to the species level with confidence. In the current study, the WW group exhibited an increased relative abundance of *H. pullorum, F. prausnitzii, B. viscericola,* and *R. torques.* Although some researchers suggested that *H. pullorum* may be an opportunistic pathogen (23), the role of *H. pullorum* in poultry and its pathogenicity to human and chickens remains unclear. In fact, to date, limited evidence has shown that *H. pullorum* is directly linked to either human or poultry diseases (24, 25). Therefore, the role of *H. pullorum* in the chicken gut needs to be further studied. Similar to the current study, a previous study showed that the prevalence of *Faecalibacterium* increased in chicken ceca from the group treated by recycled litter compared with the fresh litter group, which was concluded as a beneficial effect of recycled litter (26). As a commensal member also found to be enriched in conventionally raised chickens (27), *F. prausnitzii* was identified as a promoter of epithelial health for its ability to produce metabolites such as butyrate (28). Moreover, it was found to show anti-inflammatory activity in broilers (29). *B. viscericola* has also been characterized as an important commensal in barn-raised chickens (30–32). In addition, members from *Ruminococcus* has been characterized as important butyrate producers in the gut that degrades mucins (28, 33), however, there is limited information available concerning the role of *R. torques* in the chicken gut.

The current study unveiled alterations in a range of microbial metabolic pathways within the chicken ceca as a result of barn cleaning practices. For example, at D7, the stringent response pathway (PPGPPMET-PWY) was enriched in the cecal microbiome of the FD group which was primarily harbored by *E. coli.* The stringent response regulates genes involved in response to nutrient starvation or environmental stresses (34). It is important for bacterial virulence and persistence in the environment (e.g. resistance to antimicrobials) for a variety of taxa (34) and is used by *Bacteroides* to shift from growth to stasis (35). Furthermore, a previous study revealed that the stringent response can induce microbes to the viable but nonculturable state, a state which has strong tolerance to environmental stresses with minimum nutrient requirement (36). Thus, the enrichment of the stringent pathway may indicate that the chemical disinfection may select for tolerance to harsh environmental conditions.

At day 30, microbial metabolic pathways linked to SCFA production (acetate and lactate) and amino acid biosynthesis (L-methionine, L-lysine, L-isoleucine, and branched-chain amino acids) were enriched in the WW group. This suggests that the WW treatment led to an enhancement in the nutrient utilization functionality of the gut microbial community at day 30. Previously, Gong et al. reported that chickens infected by *Clostridium perfringens* had a significant inhibition of the pyruvate fermentation to acetate and lactate pathway II (PWY-5100) in the cecal micribiome, which was restored by the supplementation of probiotic *L. plantarum* (37). In the current study, PWY-5100 was mainly harbored by *H. pullorum* in both the FD and WW group, and to a lesser extent, *Lachnoclostridium* sp. *An76* and *Lactobacillus salivarius*. This observation suggests that these bacteria may potentially confer beneficial effects within the chicken gut at day 30.

Interestingly, PWY-1269 was shown to be mainly harbored by *H. pullorum* and *C. jejuni* in the chicken gut microbiome of the WW and FD, respectively. PWY-1269 encodes genes that produce acid sugar 3-deoxy-α-D-manno-2-octulosonate, which is a component of bacterial lipopolysaccharides (LPS) (38–40). We have previously reported that *C. jejuni* was decreased in the D30-WW group (13), which was consistent with the result from functional analyses in the current study as reflected by decreased functionality of *C. jejuni-*derived LPS genes in WW-cecal microbiome.

Regarding the ARG profiles in the chicken cecal microbiota, in the current study, the most abundant ARGs detected conferred resistance to tetracycline (*tetW, tet*(*44*)*, tet(W/N/W), tetQ, tetO* and *tet*(*32*)), MLS (*lnuC, ermB*), and aminoglocoside (*APH*(*3*)*-IIIa*); whereas β-lactam resistance was found in relatively low prevalence. Tetracycline-resistance genes, particularly *tetW,* were frequently detected in environments related to livestock farming (41–45) as well as bacteria isolated from the chicken gut (46). Previously, Munk et al. reported that the majority of ARGs in chicken fecal samples collected from multiple European countries were tetracycline-, aminoglycoside-, and MLS-resistant genes (47). Similarly, chicken fecal samples collected from China were high in aminoglycoside, tetracycline, MLS, and β-lactam resistance (48). The low abundance of β-lactam resistant genes observed in this study may link to the prohibition of prophylactic use of β-lactam in poultry farming since 2018 (49), whereas β-lactam was reported to be one of the most commonly used antibiotic in poultry production in China (50).

Regarding the ARG profiles of the chicken cecal microbiota, our observations indicate that both barn cleaning methods and the age of the chickens influenced the microbial resistome. Age emerged as the primary factor driving alterations in the resistome. In line with previous studies (51, 52), we observed that the barn-cleaning effects on the gut resistome decreased with increasing age (51, 52).

While our initial expectation was that disinfection might enhance the selection of antibiotic resistance genes (ARG), our results indicate that disinfection is linked to a reduced abundance of ARG. This finding suggests that the impact of disinfection on ARG transmission between flocks merits further evaluation as a potential strategy for controlling ARG spread. It has been reported that poultry litter harbored high densities of AMR bacteria and antibiotic residuals (53–55). Although poultry litter was removed in the current study, without chemical disinfectants, WW might preserve bacteria carrying ARGs. Therefore, it is reasonable to assume that compared to WW, the chemical disinfectants used in the current study may be more effective in controlling ARGs. Notably, at both D7 and D30, ARGs from the *erm* gene family and the glycopeptide resistant gene cluster (*vanR*) were depleted by the disinfection. Currently, more than 30 *erm* genes have been characterized, and a number of them (e.g. *ermB, ermC, ermG, ermF, ermX*) were frequently detected in livestock farming related environments (56–58). Frequent horizontal transfer of *erm* genes through mobile genetic elements has been reported within the gut microbiota (59, 60) as well as between intestinal and environmental bacteria (61, 62), accounting for their distribution among diverse taxa. Additionally, *erm* genes were found to persist stably both in the gut and in the environment 2-3 years after the removal of antibiotic selection pressure (58, 63). Furthermore, *erm* genes were also known to spread through poultry dust (45, 64), indicating possibilities of persistent *erm* residues from the previous production cycles. The *vanS/vanR -* two-component regulatory system is important in activating and regulating transcription of the glycopeptide gene cluster. Vancomycin resistant genes have been detected in poultry farms (65) and products (66). Similar to *erm* genes, a recent study revealed a glycopeptide resistant gene cluster persisting in the environment for 20 years (67). In addition, glycopeptide resistance gene clusters are also highly transferable via plasmids (66, 68), making it difficult to identify the main carriers.

In the present study, we found that the relative abundance of total ARGs was higher in the ceca of D7 chickens compared to D30. Previously, Lebeaux et al reported similar results in human infants showing that the overall relative abundance of ARGs was higher at 6 weeks than 1 year (69). We noted a heightened proportion of genes encoding antibiotic target alteration coupled with a diminished proportion of genes encoding antibiotic efflux in the chicken gut resistome from day 7 to day 30. The successional change of the chicken gut microbiota may have contributed to the alteration of cecal ARG composition. LEfSe analysis suggested that the family *Lachnospiraceae* and *Enterobacteriaceae* were strong biomarkers of the chicken cecal microbiota at D7. In addition, spearman correlation revealed that the relative abundance of the MFS antibiotic efflux pump family was positively correlated to *Lachnospiraceae* (*Lachnoclostridium* sp *An76*), further supporting the role *Lachnospiraceae* may play in the gut microbial resistome (Figure S2). Juricova et al. compared ARG sequences and bacterial genomes and reported that the family *Lachnospiraceae* was an important reservoirs for MFS antibiotic efflux pumps (46). Thus, the predominance of the MFS antibiotic efflux pumps detected in the D7-cecal resistome may be partially explained by the predominance of *Lachnospiraceae* in early life. In addition, *Enterobacteriaceae* (*E. coli* and *E. albertii*) were shown to positively correlated with the abundance of the RND antibiotic efflux pumps (Figure S2). *Enterobacteriaceae* are known to harbor the RND antibiotic efflux pumps (70, 71), and many RND genes were enriched in *Enterobacteriaceae* in chicken litter and cloacal samples (52). Thus, the enriched RND antibiotic efflux pump family at D7 may be a consequence of the early colonization of *Enterobacteriaceae*. In addition, among all taxa, *E. coli* was associated with the highest number of ARG families (Figure S2). Interestingly, in the human infant resistome study, Lebeaux et al. concluded that the human early-life resistome composition was primarily driven by *E. coli* (69).

The D30 resistome was associated with antibiotic inactivation genes, particularly genes encoding LNUs (mainly *lnuC* gene) and tetracycline inactivation enzymes (mainly *tetX* gene and its variants). Lincosamide resistant gene *lnuC* has been identified in the genus *Streptococcus* (72), and was shared extensively between different phyla (73). Emerging evidence also showed that *C. jejuni and C. coli* harbor *lnuC* (74–77). The increased relative abundance of *Streptococcaceae* and *Campylopbacteraceae* at D30 may partially explain the enriched LNU family observed in older chickens. Consistent with the effect of age, LNU genes were also more predominant in adult cattle compared to calves (51). Interestingly, *Bacillus subtilis* was positively correlated to the LNU gene family (Figure S2). *B. subtilis* has exhibited an ability to naturally activate the competence state and uptake foreign DNA (78), making it as a potential carrier of LNUs. However, the degree to which microbial compositions affect ARG composition is still unclear (79), especially in the case of highly mobile ARGs such as *lnuC*.

## Conclusion

This is the first study to report that the impact of disinfectants in broiler production on microbial functional capacity and resistome. We showed that the barn chemical disinfection may alter the composition of the chicken gut microbiota and thereby lead to decreased microbial functional capacity for amino acid and SCFA metabolism. Conversely, although differences were modest, FD may be beneficial through lowering abundance and diversity of ARGs.

